# Genomic structural equation modeling of impulsivity and risk-taking traits reveals three latent factors distinctly associated with brain structure and development

**DOI:** 10.1101/2025.06.05.657840

**Authors:** Mari Shishikura, Lang Liu, Eric Yu, Rui Zhu, Laura Vilar-Ribó, Renata B Cupertino, Abraham A. Palmer, Sandra Sanchez-Roige, Uku Vainik, Filip Morys, Ziv Gan-Or, Bratislav Misic, Alain Dagher

## Abstract

**Background:** Impulsivity is a multifaceted transdiagnostic trait that emerges in childhood. Research has identified genetic loci and brain systems associated with different facets of impulsivity and risk-taking. However, how these genetic underpinnings overlap across different facets, and how they are associated with brain development during childhood remain unknown.

**Methods:** Using genomic structural equation modeling on 17 impulsivity and risk-taking traits, we identified latent factors capturing overlapping genetic architecture. We then calculated polygenic scores for these factors using Adolescent Brain Cognitive Development Study data (N = 4,142) and examined their associations with brain structure, development, and behavior in children aged 9-14 years. Finally, we tested whether socioeconomic status modulated the associations between latent polygenic scores and brain structures.

**Results:** We identified three distinct genetic latent factors, which we label lack of self-control, reward drive, and sensation seeking. In children, polygenic scores for the three factors showed associations with distinct brain patterns: lack of self-control associated with reduced prefrontal cortical thickness, reward drive with increased subcortical cellular density, and sensation seeking with increased cortical surface area and white matter integrity. Longitudinally, lack of self-control predicted slower white matter development. The association between polygenetic score for lack of self-control and white matter mean diffusivity was modulated by socioeconomic status.

**Conclusions:** We identified three genetically distinct dimensions of impulsivity and risk-taking with separable neurodevelopmental origins. These genetic predispositions manifested as distinct brain patterns as early as ages 9–10. Environmental experience modulated some of the genetic effects on brain development.

## Introduction

Impulsivity refers to acting quickly without sufficient consideration for consequences (1,2) and can also encompass the tendency to take risks (3,4). Impulsivity is a transdiagnostic trait observed in many psychiatric disorders, including attention-deficit/hyperactivity disorder (5,6), substance use disorders (7–9), bipolar disorder (10–12), and schizophrenia (13,14).

Impulsivity is typically assessed using self-reported questionnaires, such as the Barratt Impulsiveness Scale (BIS-11) (15) and the UPPS-P (16,17). Alternatively, laboratory tasks, such as the Delay Discounting task, can be used to measure one’s preference for smaller immediate rewards over larger delayed rewards (18). In addition, risky behaviors—such as speeding, alcohol consumption, and smoking—are often evaluated alongside impulsivity, as they reflect a tendency to engage in activities with potential negative health consequences (19).

Impulsivity is therefore a multifaceted construct (20–27), with several theories proposed to characterize its underlying dimensions. For example, dual-process models identify two competing systems: one for cognitive control, which regulates imprudent impulses, and one for motivational drive, which promotes reward or sensation seeking (28–32).

Neuroimaging studies mirror the multidimensional nature of impulsivity, identifying multiple brain structures and networks associated with facets of impulsivity. Trait impulsivity has been associated with lower gray matter volume in the prefrontal cortex (33). Meanwhile, task-based functional MRI research found that impulsive decision-making is associated with greater activation of ventromedial prefrontal cortex and ventral striatum, while inhibition of impulsive responses recruits regions in fronto-parietal control networks (34,35).

Twin studies have shown that impulsivity and risk-taking traits are heritable, with heritability estimates varying between 30 and 60% (36–42). Genome-wide association studies (GWASs) have further identified key loci associated with different impulsivity traits near genes implicated in neurobiological processes (43–46). However, how these genetic predispositions manifest in the brain during development is largely unknown.

Identifying the mechanisms underpinning impulsivity and risk-taking in early adolescence is of interest, as this period is marked by heightened impulsivity and increased engagement in risky behaviors (47–49). Moreover, since most individuals have not yet experienced prolonged substance use during early adolescence (50–52), this stage offers a unique opportunity to disentangle the brain mechanisms driving impulsivity and risk-taking from the effects of substance exposure.

The aim of the present study was three-fold. First, we aimed to identify genetic constructs underlying different facets of impulsivity and risk-taking by applying genomic structural equation modeling to 17 relevant traits from published GWASs (44–46,53,54). Although the genetic architecture of impulsivity has been previously explored (55–58), we extended this work by jointly analyzing risk-taking and impulsivity traits using a data-driven framework. Second, we examined whether the genetic predisposition for impulsivity and risk-taking is associated with brain structures and cognitive function during early development. To this end, we utilized data from the Adolescent Brain and Cognitive Development ^SM^ (ABCD) Study, the largest genetic and neuroimaging dataset of youth to date. Finally, since socioeconomic status (SES) has been linked to both impulsivity and brain development (59–62), we examined how genetic predisposition for impulsivity and SES factors such as parental education and household income jointly contribute to differences in brain structures among children and adolescents.

## Materials and Methods

### GWAS of impulsivity and risk-taking behavior

We used published GWAS summary statistics of 17 traits related to impulsivity and risk-taking behaviors (Table. 1). We included Body Mass Index (BMI) and Education Attainment (EA) for the initial analyses, given their relevance to impulsivity (63–68). All GWASs were derived from individuals of genetically inferred European ancestry.

### Genomic Structural Equation Modeling (genomic SEM)

*Genomic SEM* R package v0.0.5 (71) was used to build and test factor models, following our previous work (55–58,72). While previous hypothesis-driven models successfully captured aspects of the genetic architecture of impulsivity, here we adopted a data-driven (hypothesis-free) approach (73) to further explore how impulsivity and risk-taking traits are organized jointly (See Supplementary Information for detailed descriptions). Model fit was evaluated using comparative fit indices (CFI) and standardized root mean square residuals (SRMR).

### Multivariate GWAS

*Genomic SEM* R package (71) was used to conduct multivariate GWAS, where SNP associations with the latent factors derived from genomic modeling were estimated. The effective sample size was calculated as described by Mallard et al. (74). For subsequent analyses, we omitted SNPs that had Q_SNP_ p-values below 5e-8 (See Supplementary Information).

The SNP2GENE function from the Functional Mapping and Annotation (FUMA) v1.5.2 pipeline (75) was used to identify genomic risk loci and lead SNPs.

### Gene-based analyses

We used MAGMA v1.08 (76) to conduct gene association, gene-set enrichment, and gene property analyses for each latent factor. These analyses tested whether GWAS-implicated genes were preferentially associated with specific gene-sets, expression in particular tissues, or expression during specific developmental epochs. (See Supplementary Information for reference datasets. For the developmental epoch analysis, arthritis GWAS (77) was used as a negative control to support that associations reflected impulsivity-specific patterns, not general developmental expression trends.

### ABCD dataset

#### Participants

To examine how the genetic predisposition of impulsivity and risk-taking unfolds during development, we used data from the ABCD Study ®. This is a longitudinal cohort study, with genetic, demographic, behavioral, and brain-imaging data collected across 21 sites in the US (78–80). We used neuroimaging and genetic data from the ABCD 5.1 release. The participants were recruited at ages 9 to 10 and undergo follow-up MRI scans every 2 years. We included participants of genetically inferred European ancestry to match the population of the GWASs. This totaled 4,142 participants at baseline (mean age: 119.4 months, females: 1,958), 3,260 participants at 2-year-follow up (mean age: 143.6 months, females: 1,451), and 1,346 participants at 4-year-followup (mean age: 169.2 months, females: 606). Subjects with at least two measurements were included in the longitudinal analysis (N= 3,552, females: 1591).

#### Neuroimaging data

Neuroimaging data was preprocessed by the ABCD consortium as described elsewhere (81). We used cortical surface area and thickness measures derived from Desikan-Killiany (DK) Atlas (82), and white matter (WM) tracts labeled using AtlasTrack (83). Fractional anisotropy (FA) and mean diffusivity (MD) were obtained for each white matter tract. We also included hindered normalized isotropic component (HNI), representing extracellular diffusion, and restricted normalized isotropic component (RNI), representing intracellular diffusion, for subcortical structures (81,84,85). We harmonized imaging measures using longComBat to account for bias arising from different scanning sites (86). Imaging data that did not pass the ABCD recommended QC was excluded (81). In total, we analyzed 234 regional brain measures: cortical surface area and thickness (68 regions each), subcortical volume (16 structures), subcortical HNI and RNI (14 each), and FA/MD across 27 white matter tracts.

#### Genetic data

To derive individual-level genetic predisposition for impulsivity and risk-taking (see below), we analyzed the genotype data from ABCD participants. These were obtained using the Affymetrix Axiom Smokescreen Array platform (87,88). Individuals with a high rate of genotype missingness (0.05), high or low rate of heterozygosity (F-statistic <−0.15 or >0.15), mismatched sex labeling, and high relatedness (π>0.125) were excluded from the sample. We also removed data from genotyping batch 461. Ancestry outliers were detected using HapMap3 data in R v4.0.1. See Supplementary Information for detailed descriptions.

#### Behavioral and Cognitive data

To examine whether genetic predisposition captured relevant phenotypes, we looked into impulsivity-related measures in the ABCD dataset. These included the UPPS-P Impulsive Behavior Scale for children (80) and the Delay Discounting task (89–91). We also included the Behavioral Inhibition Scale and Behavioral Activation Scale (BIS/BAS), which assess sensitivity to punishment and reward (80,92), and the Externalizing and Internalizing Behavior Composite Score from the Child Behavior Checklist (CBCL), completed by parents (80). Lastly, we included BMI in our analyses, given its well-documented association with impulsivity (63–68).

We further explored associations with broader domains of cognitive function. We included the Little Man Task (visuospatial processing) (89), Wills Problem Solving Task (problem-solving skills) (93–95), Matrix Reasoning from the Wechsler Intelligence Scale for Children-V (nonverbal reasoning) (89), and the Emotional Stroop Task (96). Additionally, we calculated a composite executive function score based on NIH Toolbox tasks, following previous studies (89,97–100).

Most measures in our analysis come from the baseline data collection, except the Delay Discounting Task and Wills Problem Solving Task, which were from the 1-year follow-up.

### Socioeconomic Status (SES) of Parents

We assessed socioeconomic status using average parental education (in years), household income, marital status, and the Area Deprivation Index (ADI)—a composite measure of neighborhood disadvantage based on 17 socioeconomic indicators, where higher scores indicate greater deprivation (101,102).

#### Polygenic Score (PGS) calculation

PGSs for the latent factors identified in genomic SEM were calculated using the PRS-CS pipeline (103), accounting for linkage disequilibrium (LD) using 1000 Genomes Phase 3 reference data.

#### Longitudinal modeling of neuroimaging data

To assess the developmental trajectory of brain structures, we constructed a latent growth curve model (LGCM) (104,105) for each brain measure (5 global measures and 234 regional measures), using *lavaan* package (106) in R (107). Each model estimated group-level intercepts and slopes, as well as individual trajectories. Percentage change per annum was calculated as slope/intercept×100. All models had a good fit with CFIs above 0.90 and SRMRs below 0.10.

#### Partial Least Square

We used Partial Least Squares (PLS) analysis to identify latent patterns of brain structure associated with latent factor PGSs in a multivariate framework. PLS decomposes the covariance between two datasets—in this case, PGS value and brain features—into latent variables that maximally capture their shared variance (108–110). Before analysis, both datasets were z-scored, and brain features were residualized for age, sex, 15 genetic principal components (PCs), and genotyping batch. The significance of latent factors was tested via permutation testing. A Bootstrap Ratio (BSR) was calculated for each variable to capture both the strength and the reliability of its contribution to the latent pattern. Cross-validation was performed to assess robustness and generalizability (111,112). Detailed methods are provided in Supplementary Information.

The same PLS framework was applied to longitudinal brain change data, with the brain matrix replaced by annual percent change in brain features.

#### Interaction analysis with PGSs and SES

Linear regression models with an interaction term (PGS × SES) were constructed to test whether SES modulated the relation between PGS and global brain measurement. Each model was adjusted for age, sex, 15 genetic PCs, and genotyping batch. The interaction term was tested for significance, and false discovery rate correction was applied for multiple comparison (See Supplementary Information for detailed descriptions).

#### Dominance Analysis

We used dominance analysis to assess the relative contribution of genetic and environmental predictors to global brain measurements in separate multiple regression models (113,114). Genetic predictors included latent factor PGSs, while environmental predictors comprised parental education, household income, marital status, and ADI. Outcomes included cortical surface area, cortical thickness, subcortical volume, white matter fractional anisotropy, mean diffusivity, and subcortical cellularity (HNI, RNI).

## Results

### Genetic architecture of Impulsivity and Risk-Taking

To examine genetic relationships among impulsivity-related traits, we estimated genetic correlations across 17 GWASs—15 indexing impulsivity and risk-taking, plus BMI and educational attainment (EA) due to their established associations with impulsivity (Table 1). Most of the traits were significantly correlated (average | r_g_ | = 0.30), with stronger correlations within certain subgroups (max| r_g_ | between Drug Exposure and Smoking Initiation (r_g_ = 0.83, SE = 0.02), and lower or not significant correlations for a few others (min| r_g_ | between UPPS-P Lack of Perseverance and Sensation Seeking (r_g_ = _−_0.0013, SE = 0.04). Hierarchical clustering on the correlation matrix suggested three subgroups (Fig. 1A). EA was negatively correlated with most impulsivity traits except for positive associations with Automobile Speeding Propensity, Risk Tolerance, and Sensation Seeking, which clustered together.

**Figure 1.**
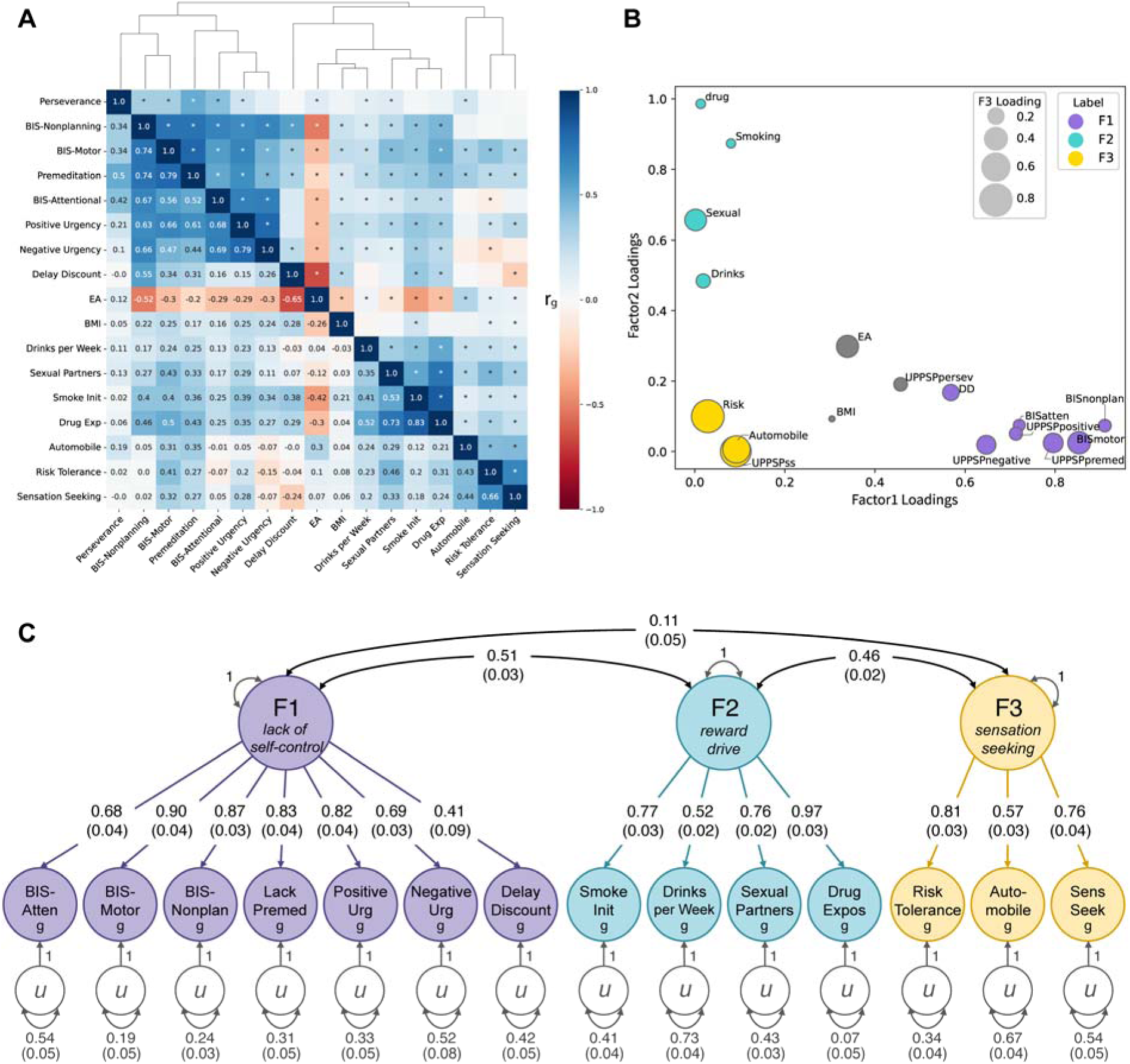
Genetic relationships among impulsivity and risk-taking behaviors. **A)** Genetic correlation matrix among 17 phenotypes of interest. **B**) Scatterplot of standardized factor loadings from explanatory factor analysis. **C**) Genomic SEM model with three latent factors. Residual covariances are omitted for visual clarity (See Fig. S1 in Supplementary Information).

**Table 1.**
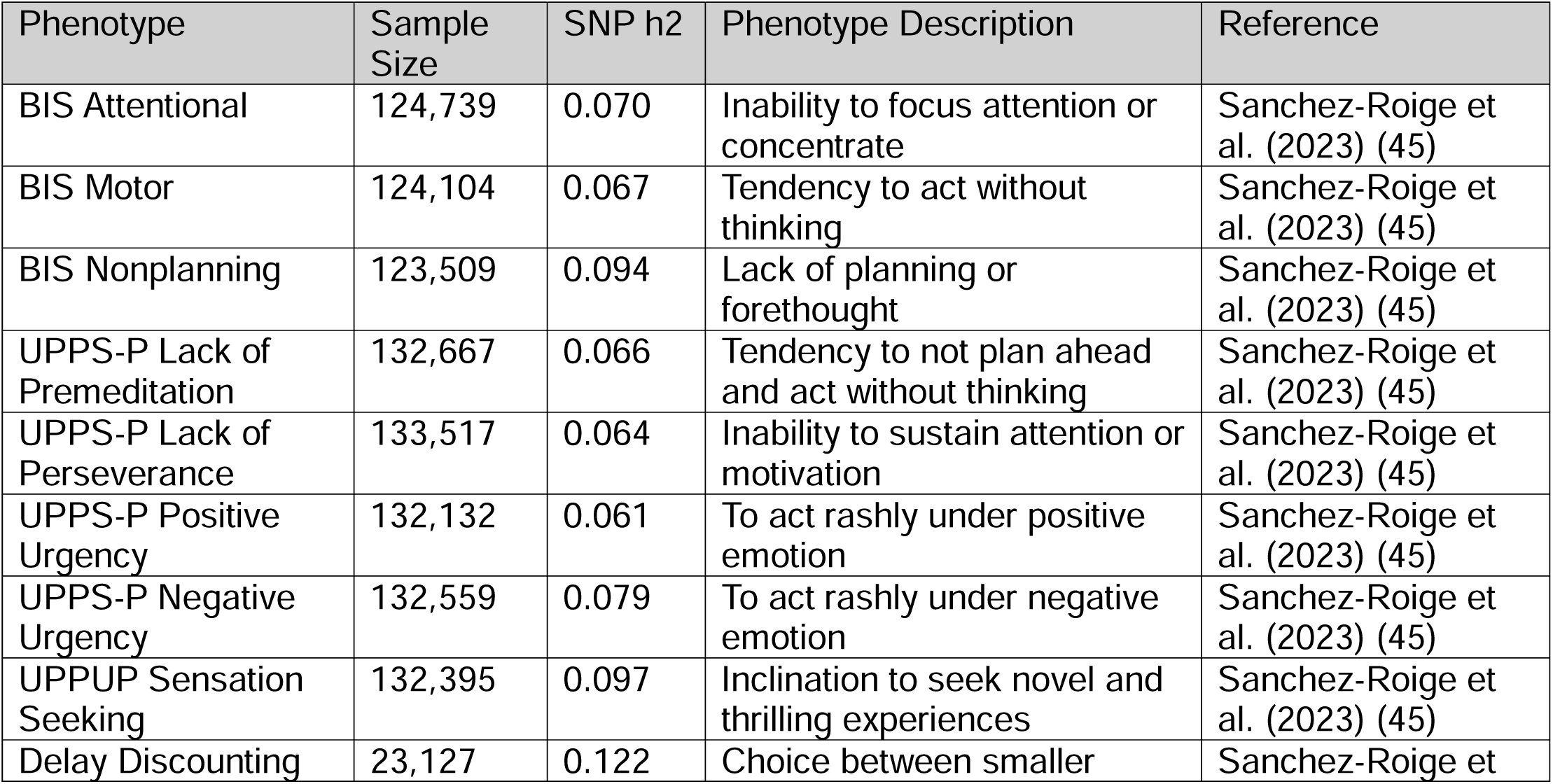
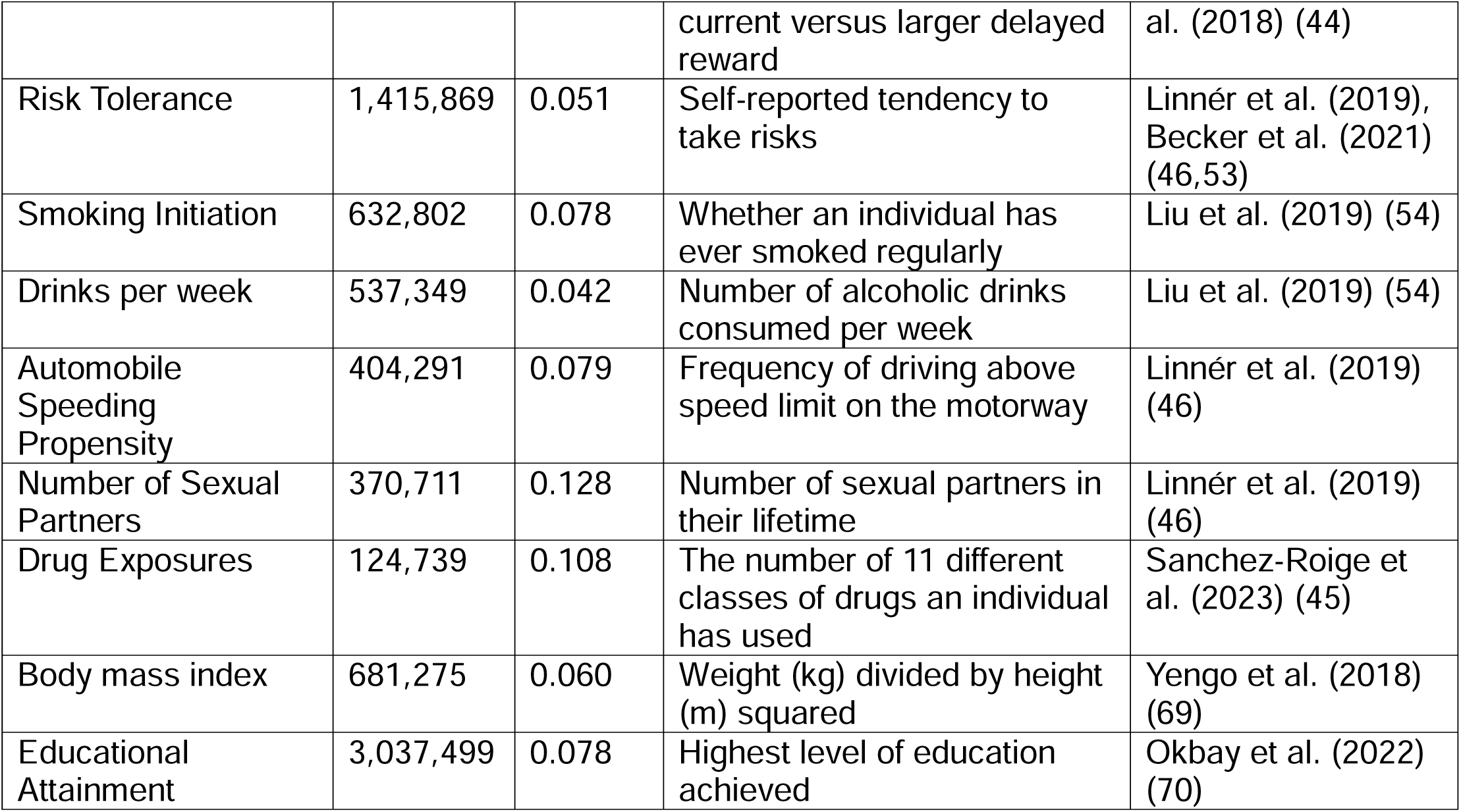
GWASs related to impulsivity and risk-taking behaviors. We used summary statistics made available by the original papers, except for Risk Tolerance and Educational Attainment. For these, we conducted meta-GWAS, following the methods described in the original paper. This is due to the limitation of data availability (See Supplementary Information). SNP h2: SNP-based heritability. None of these GWAS included individuals from the ABCD cohort.

**Table 2.** Genome-wide significant loci associated with lack of self-control, reward drive, and sensation seeking. For reward drive and sensation seeking, only top 10 most significant loci are shown. List of all significant loci (66 for *reward drive* and 45 for *sensation seeking*) are shown in Supplementary Tables S2 and S3, respectively.

| rsID | chr | pos | Non effect | effect | MAF | P value | Beta | SE | Nearest gene | function |
| --- | --- | --- | --- | --- | --- | --- | --- | --- | --- | --- |
| **Lack of self-control** |  |  |  |  |  |  |  |  |  |  |
| rs1897699 | 3 | 85489304 | T | A | 0.38 | 6.18E-10 | 0.020 | 0.0032 | *CADM2* | intronic |
| rs936146 | 7 | 114294405 | C | G | 0.41 | 1.18E-08 | -0.018 | 0.0032 | *FOXP2* | intronic |
| **Reward drive** |  |  |  |  |  |  |  |  |  |  |
| rs3802848 | 11 | 112912387 | C | A | 0.44 | 9.90E-27 | -0.021 | 0.0015 | *NCAM1* | intronic |
| rs6788098 | 3 | 85624131 | T | A | 0.37 | 2.58E-26 | 0.022 | 0.0016 | *CADM2* | intronic |
| rs1143702 | 1 | 44086831 | T | C | 0.36 | 3.52E-16 | 0.017 | 0.0016 | *PTPRF* | exonic |
| rs6493275 | 15 | 47676579 | T | A | 0.21 | 3.05E-14 | -0.018 | 0.0018 | *SEMA6D:CTD-2050N2.1* | ncRNA intronic |
| rs542883 | 2 | 45143382 | C | G | 0.42 | 4.11E-14 | -0.015 | 0.0015 | *RP11-89K21.1* | intergenic |
| rs1549212 | 5 | 166996722 | T | C | 0.36 | 8.59E-14 | 0.015 | 0.0016 | *TENM2* | intronic |
| rs1591915 | 10 | 104646849 | T | C | 0.42 | 1.03E-13 | -0.015 | 0.0015 | *C10orf32-ASMT:AS3MT* | intronic |
| rs266059 | 2 | 104084908 | G | A | 0.50 | 5.71E-13 | 0.014 | 0.0015 | *AC092568.1* | intergenic |
| rs2088088 | 6 | 98887788 | T | C | 0.34 | 1.83E-12 | -0.015 | 0.0016 | *RP11-436D23.1* | intergenic |
| rs1899896 | 8 | 93201036 | T | C | 0.31 | 5.54E-12 | -0.015 | 0.0016 | *RP11-777J24.1* | intergenic |
| **Sensation seeking** |  |  |  |  |  |  |  |  |  |  |
| rs6790699 | 3 | 85624189 | G | A | 0.37 | 4.85E-54 | 0.021 | 0.0010 | *CADM2* | intronic |
| rs4841503 | 8 | 11024326 | C | A | 0.47 | 1.53E-26 | 0.014 | 0.0001 | *XKR6* | intronic |
| rs3764002 | 12 | 108618630 | T | C | 0.26 | 2.04E-15 | 0.012 | 0.0011 | *WSCD2* | exonic |
| rs62474683 | 7 | 115020725 | G | A | 0.47 | 3.07E-15 | 0.010 | 0.0010 | *RP11-222O23.1* | intergenic |
| rs13095117 | 3 | 18745500 | G | A | 0.40 | 1.15E-14 | 0.010 | 0.0010 | *AC144521.1* | ncRNA intronic |
| rs7618871 | 3 | 136400420 | T | G | 0.43 | 1.47E-12 | 0.009 | 0.0010 | *STAG1* | intronic |
| rs12413688 | 10 | 99095544 | G | A | 0.37 | 1.87E-12 | 0.009 | 0.0010 | *RP11-452K12.4* | ncRNA intronic |
| rs4551267 | 7 | 127175236 | T | C | 0.45 | 2.51E-12 | 0.009 | 0.0010 | *GCC1* | intergenic |
| rs1015511 | 7 | 114142250 | G | A | 0.48 | 2.56E-12 | -0.009 | 0.0010 | *FOXP2* | intronic |
| rs556560 | 5 | 102623924 | G | A | 0.34 | 5.10E-12 | -0.009 | 0.0010 | *C5orf30* | intergenic |

We then modeled genetic covariances of all traits using Genomic SEM. We determined that a three factor model best explained the genetic covariance (Fig. 1B, 1C; CFI=0.907, SRMR=0.088).

The first latent factor appeared to capture *lack of self-control*. This included Barratt Impulsiveness Scale subscales, UPPS-P Premeditation, Positive Urgency, Negative Urgency, and Delay Discounting. The second factor can be labeled as *reward drive*, including Ever smoker, Drinks per week, Number of sexual partners, and Drug Exposure. Finally, the third factor can be conceptualized as *sensation seeking* tendency, encompassing Risk Tolerance, UPPS-P Sensation Seeking, and Automobile Speeding Propensity.

### Multivariate GWAS

Multivariate GWAS identified 2, 66, and 45 genomic risk loci for *lack of self-control*, *reward drive*, and *sensation seeking*, respectively (Table. 2; Fig.2; Table. S1-3). Almost all loci were previously reported in the individual GWASs (Table. S4-6), except for one locus in *reward drive* near *TNRC6B* (rs139911, *p* = 1.23×10^-8^) and 6 loci in *sensation seeking* near genes *SCARNA11* (rs61183738, *p* = 4.53×10^-8^), *AGBL4* (rs3121532, *p* = 2.53×10^-8^), *ASH1L* (rs12041534, *p* = 1.15×10^-8^), *RP11-328D5.1* (rs1204678, *p* = 2.16×10^-8^), *AC079135.1:ASB18* (rs10929169, *p* = 4.05×10^-9^), and *CCDC171* (rs9298741, *p* = 1.77×10^-8^). Among these, loci with hit SNPs rs3121532 and rs1204678 have not been implicated in previous GWASs, while others are implicated in BMI, EA, or Schizophrenia (Table. S4-6).

**Figure 2.**
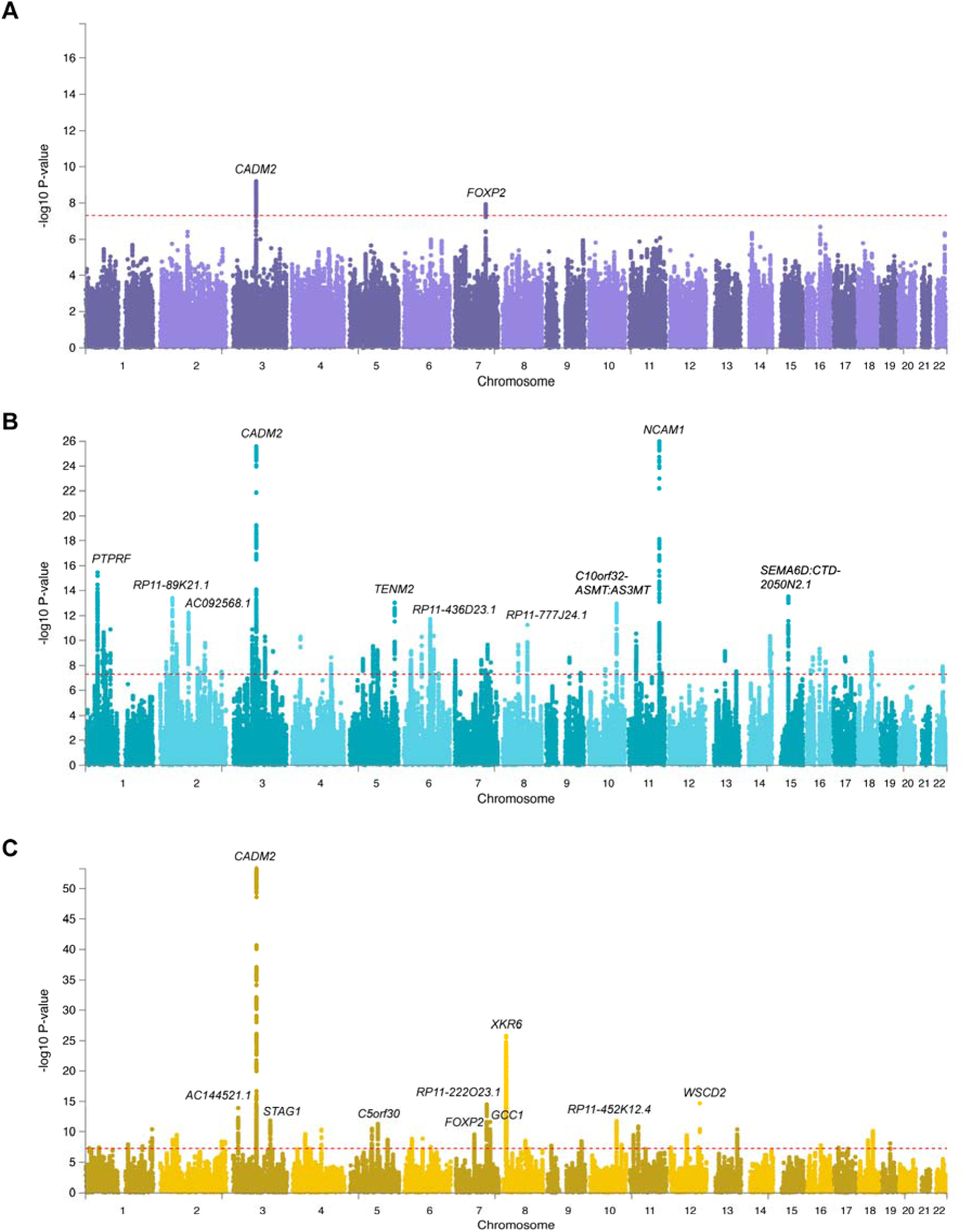
Multivariate GWAS results for the three latent factors. **A-C**) Manhattan plots for *lack of self-control* **(A)**, *reward drive* **(B)**, and *sensation seeking* **(C)**. The nearest genes for the lead SNPs are labelled (top 10 significant loci for *reward drive* and *sensation seeking*).

All three factors showed associations with loci near *CADM2* and *FOXP2*. SNPs near *CADM2* and *FOXP2* genes have been robustly implicated in previous impulsivity and risk-taking GWASs (45,46,115,116). Only 6 associations were shared between *reward drive* and *sensation seeking* (Table S2,3).

### Gene-based analyses

MAGMA identified 2, 150, and 139 genes associated with *lack of self-control*, *reward drive*, and *sensation seeking*, respectively (Table. S7-9). *CADM2* and *FOXP2* were associated with all three factors, consistent with the GWAS results, while an additional 129 and 115 genes were associated with *reward drive* and *sensation seeking*, beyond the GWAS implicated loci.

*Reward drive* was enriched for genes linked to neuronal development, while *sensation seeking* was enriched for synaptic properties (Table. S10-12). No significant enrichment was found for *lack of self-control* after multiple comparison corrections.

Tissue-specific analysis showed broad enrichment in brain tissues across all three factors (Fig. S2). Finally, BrainSpan-based gene property analyses revealed that *reward drive* was significantly enriched for genes preferentially expressed during early, early-mid, and late-mid prenatal period (*p* = 1.17×10^-3^, *p* = 2.23×10^-5^, *p* = 1.14×10^-4^, respectively), while *sensation seeking* was enriched for genes expressed in late infancy (*p* = 1.19×10^-4^, Fig. 3).

**Figure 3.**
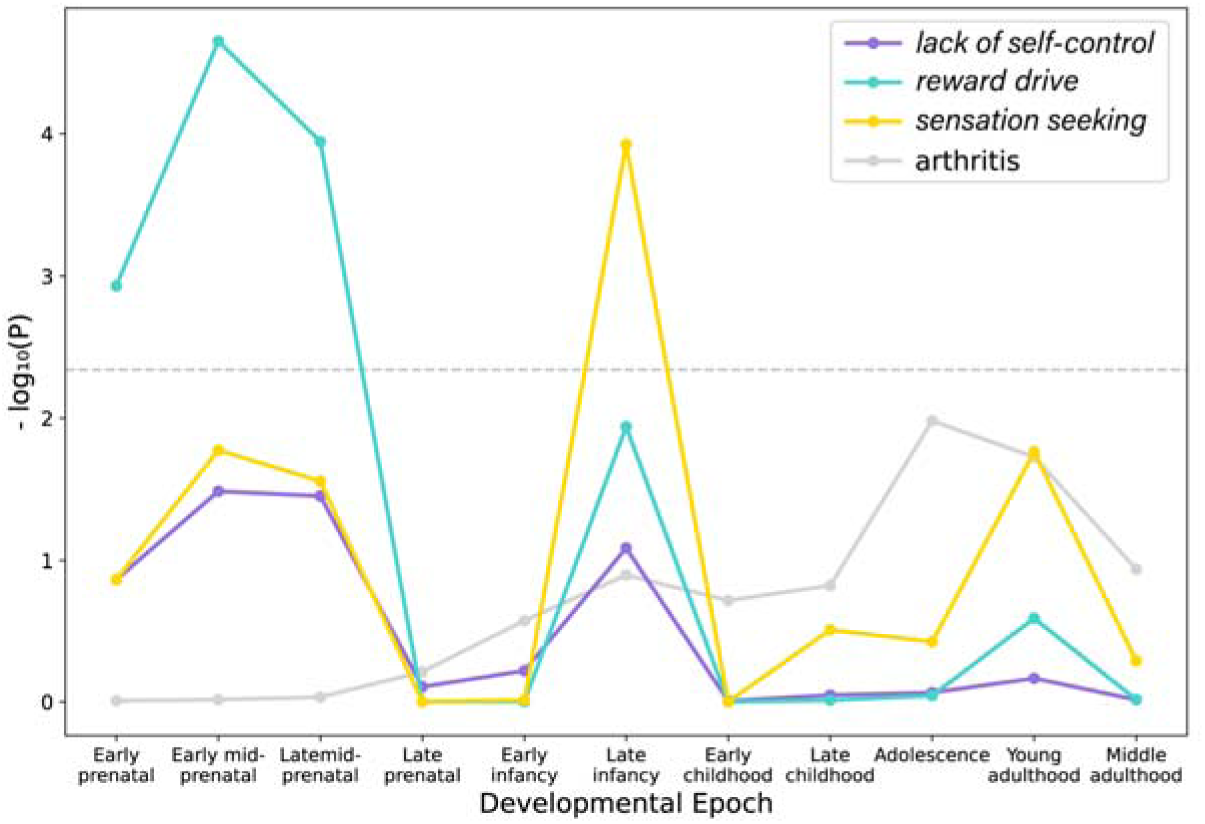
Gene expression enrichment for each developmental epoch. The y-axis refers to the significance on a -log_10_ scale and the horizontal dashed line denotes significance after multiple comparison correction. Arthritis is shown as a control.

### Polygenic analyses of impulsivity and risk-taking in children and adolescents

#### Behavioral Phenotypes

PGS for *lack of self-control*, *reward drive*, and *sensation seeking* were all generally associated with impulsivity measures and risky behaviors, although they diverged in specific association patterns (Fig. 4, Table. S13). *Lack of self-control* PGS showed significant associations with Delay Discounting (β = 0.031, *p*_FDR_ = 0.045) and UPPS-P subscales except Sensation Seeking (standardized β = 0.038−0.063). *Reward drive* PGS showed the largest association with Externalizing Behavior Scores (β = 0.096, *p*_FDR_ = 9.45×10^-10^) and was associated with all UPPS-P subscales (/3 range between 0.050 and 0.074). *Sensation seeking* PGS was significantly associated with only Sensation Seeking (β = 0.075, *p*_FDR_ = 3.33×10^-6^) and Lack of Planning (β = 0.061, *p*_FDR_ = 1.22×10^-4^) among UPPS-P subscales. Furthermore, we observed associations beyond impulsivity and risk-taking traits. All three PGSs were negatively associated with the Wills Problem Solving Scale, which captures cognitive resilience in problem solving (β = −0.053, *p*_FDR_ = 7.88×10^-4^, β = −0.062, *p*_FDR_ = 6.55×10^-5^, β = −0.032, *p*_FDR_ = 0.0453, respectively).

**Figure 4.**
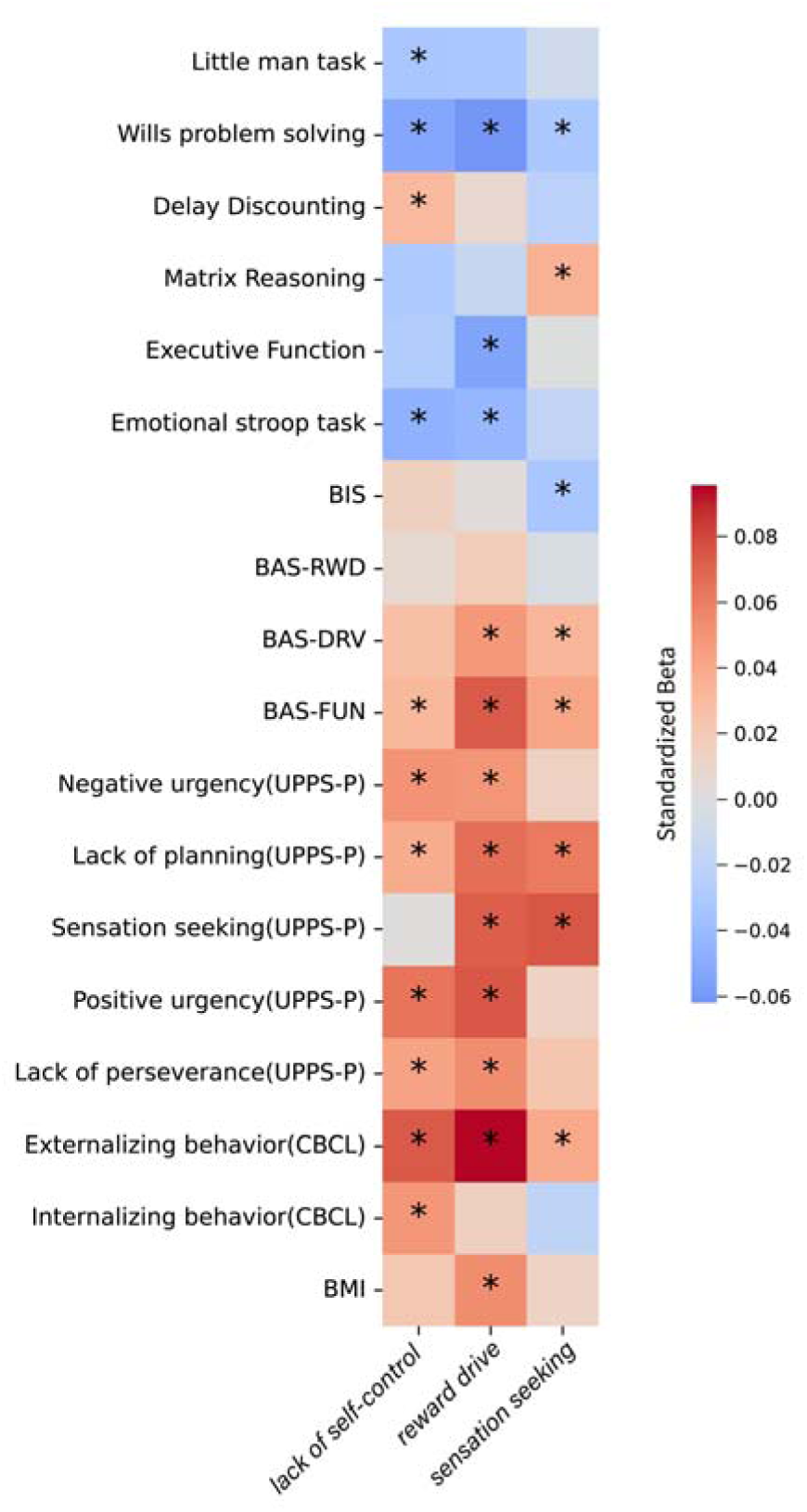
Association between PGS for lack of self-control, reward drive, and sensation seeking and behavioral and cognitive phenotypes in the ABCD cohort. Age, sex, 15 genetic PCs, and genotyping batch were corrected for all linear regression analyses. Asterisk denotes significance after FDR correction. Measures are from baseline collection (age 9-10) except for Delay Discounting and Wills problem solving (from 1 year follow-up, age 10-11). BIS: Behavioral Inhibition Scale, BAS: Behavioral Activation Scale, RWD: Reward Responsiveness, DRV: Drive, FUN: Fun Seeking.

#### Brain Measures

Using PLS analysis, we examined whether each PGS was associated with a distinct pattern of brain structural characteristics in the ABCD dataset.

*Lack of self-control* PGS was significantly associated with a latent brain pattern (*p*_perm_ = 0.010; Fig. 5A), characterized by reduced cortical thickness of right hemisphere precentral cortex (BSR = −4.01), left cuneus (BSR = −3.44) and insula (BSR = −3.18), and reduced surface area of left insula (BSR = −3.62) and right pars orbitalis (BSR = −3.03; Table. S14).

**Figure 5.**
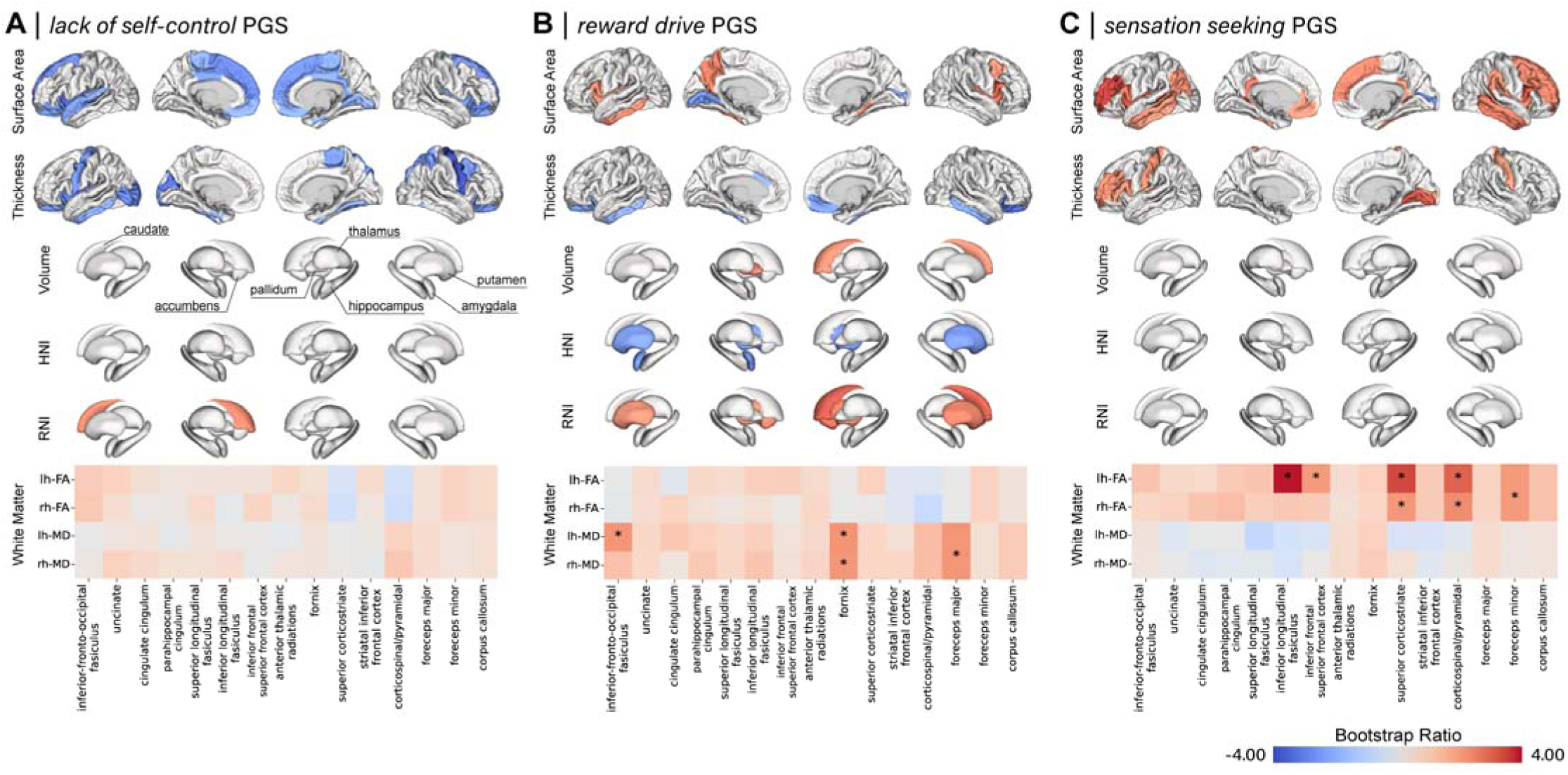
Brain patterns associated with genetic predisposition for impulsivity and risk taking. Bootstrap ratios from the PLS analyses between brain regional measures and **A**) *lack of self-control* PGS, **b**) *reward drive* PGS, and **c**) *sensation seeking* PGS. Regions with Bootstrap Ratio (BSR) values exceeding the threshold corresponding to p < 0.05 are highlighted, indicating significant contribution of the brain measures to the associations. HNI: hindered normalized isotropic component, RNI: restricted normalized isotropic component, FA: fractional anisotropy, MD: mean diffusivity, lh: left hemisphere, rh: right hemisphere.

*Reward* drive PGS and brain measurements also showed a significant association (*p*_perm_ = 0.040; Fig. 5B, Table. S15). The identified brain pattern was characterized by thinner lateral orbitofrontal cortex (BSR_right_ = −3.42, BSR_left_ = −2.32), larger surface area in pars opercularis (BSR_right_ = 2.97, BSR_left_ = 2.26) and insula (BSR_right_ = 2.79, BSR_left_ = 2.18), and higher RNI, i.e., higher cellular density (117,118), in right caudate (BSR = 3.13) and accumbens (BSR_right_ = 2.91, BSR_left_ = 2.29).

Finally, *sensation seeking* PGS also significantly associated with a latent brain pattern (*p*_perm_ = 0.014; Fig. 5C, Table. S16). The brain pattern was characterized by higher FA in left inferior longitudinal fasciculus (BSR = 3.85), superior corticostriate (BSR_left_ = 3.32, BSR_right_ = 2.02), and corticospinal tracts (BSR_left_ = 2.95, BSR_right_ = 2.24), and larger surface area in rostral middle frontal cortex (BSR_left_ = 3.49, BSR_right_ = 2.49), insula (BSR_left_ = 3.25, BSR_right_ = 2.40), and pars opercularis (BSR_left_ = 3.25, BSR_right_ = 2.63).

We ran cross-validation to confirm the generalizability of the PLS analyses. The resulting mean out-of-sample brain-PGS correlation was significant for all three analyses (*lack of self-*control PGS: r(1028) = 0.042, *p*_perm_ = 0.0198; *reward drive* PGS: r(1028) = 0.050, *p*_perm_ = 0.0346; *sensation seeking* PGS: r(1028) = 0.039, *p*_perm_ = 0.0099).

#### Brain Development

We also tested whether the PGSs were related to the brain developmental trajectory. *Lack of self-control* PGS showed a significant association with changes in brain structure (*p*_perm_ = 0.020; Table. S17). As each measurement has a distinct direction of change during this period (Fig. 6A), we translated the BSR to the level of development (Fig. 6B).

**Figure 6.**
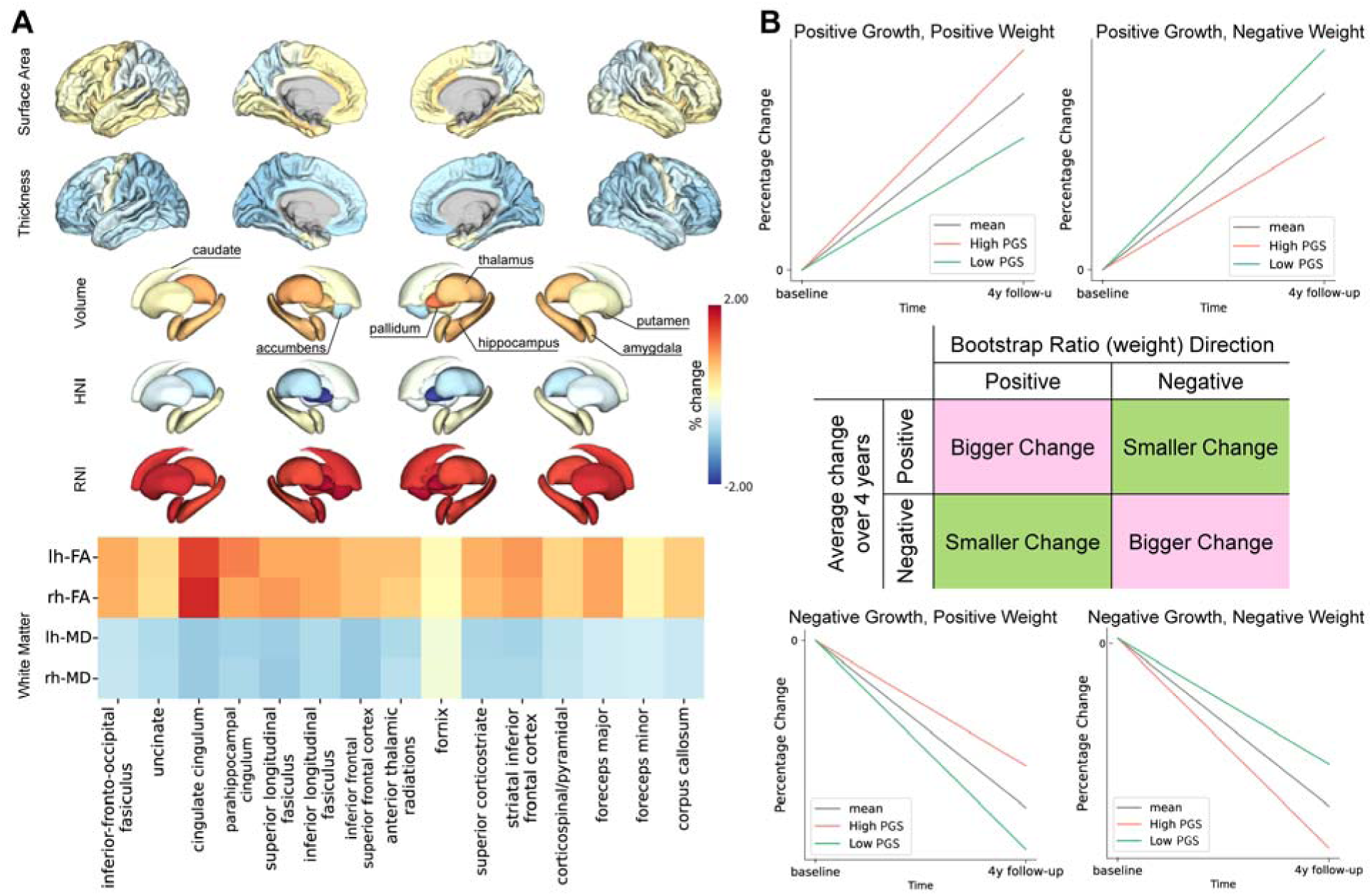
Brain development trajectory and the directional interpretation of PLS. **A)** Annual percentage change for regional brain measurements derived from latent growth curve models. **B**) Schematic panel of how developmental direction and Bootstrap ratio (BSR) translates to levels of development. A positive BSR in regions with positive growth (top left), or a negative BSR in regions with negative change (bottom right) indicates that higher PGS is associated with bigger change (accelerated development). Conversely, a negative BSR in regions with increasing values (top right), or a positive BSR in regions with negative developmental change (bottom left) indicates that having higher PGS is associated with smaller change (slowed development). HNI: hindered normalized isotropic component, RNI: restricted normalized isotropic component, FA: fractional anisotropy, MD: mean diffusivity, lh: left hemisphere, rh: right hemisphere.

The association with *lack of self-control* PGS was characterized by smaller increase in WM FA values—most prominent in the right inferior frontal superior frontal tract (BSR = −3.40), left uncinate (BSR = −3.24), and cingulate cingulum (BSR_left_ = −3.17, BSR_right_ = −3.05)—as well as smaller decrease in HNI of right amygdala (BSR = 2.22), putamen (BSR = 2.06), and left hypothalamus (BSR = 2.10; Fig. 7, Table S14). It also involved greater cortical thinning in left pericalcarine (BSR = −3.22) and more shrinking of right lateral occipital cortex (BSR = −2.98) and pars opercularis (BSR = −2.75). Cross-validation confirmed generalizability (r(785) = 0.036, *p*_perm_ = 0.0396).

**Figure 7.**
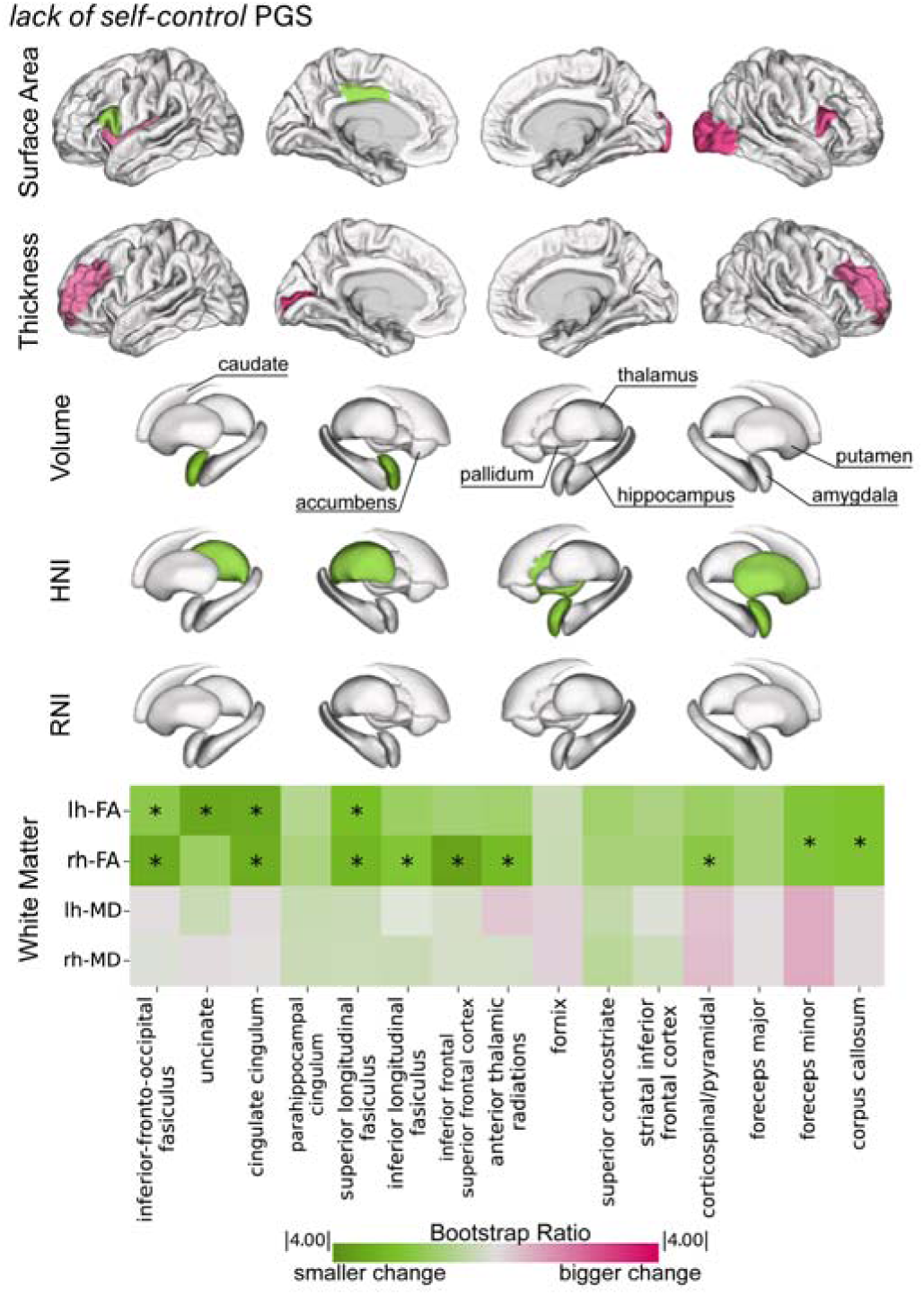
Brain developmental patterns associated with genetic predisposition for lack of self-control. Bootstrap ratios (BSRs) from PLS analysis between regional developmental trajectories and *lack of self-control* PGS. The figure is color-coded to reflect whether *lack of self-control* PGS is associated with bigger growth (pink) or smaller growth (green). BSR values exceeding the threshold corresponding to *p* < 0.05 are highlighted. HNI: hindered normalized isotropic component, RNI: restricted normalized isotropic component, FA: fractional anisotropy, MD: mean diffusivity, lh: left hemisphere, rh: right hemisphere.

We found no significant association between *reward drive* or *sensation seeking* PGS and developmental changes in brain measurements.

### Relation with Socioeconomic status

#### Interaction Analysis

We explored whether the effect of the PGS depended on socioeconomic status. The association between *lack of self-control* PGS and overall WM MD was moderated by ADI (β = 0.057, *p*_FDR_ = 0.039; Fig. 8A). Further inspection revealed that individuals with higher ADI had a positive association between PGS and WM MD, whereas those with lower ADI showed a negative association (Fig. 8B).

**Figure 8.**
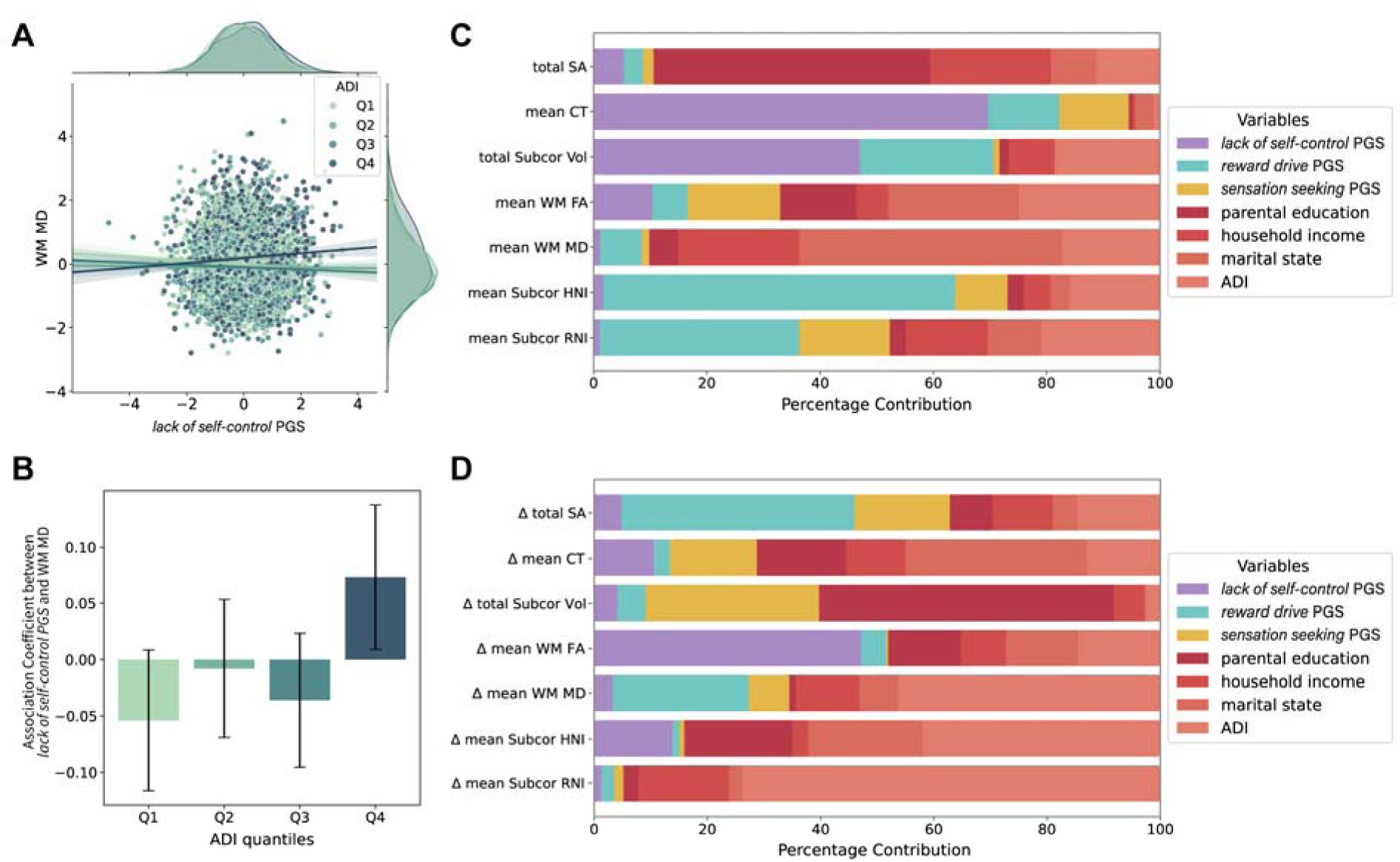
Interacting and collective effect of genetic predisposition and socioeconomic status on the brain. **A)** Scatterplot of association between *lack of self-control* PGS and WM MD, in relation to ADI quantile. Each point is color-coded according to the ADI quantile to which it belongs. The regression line represents the association for each quantile. **B**) Bar plot showing the association coefficient between *lack of self-control* PGS and WM MD at different ADI quantiles. The y axis shows the standardized beta estimates from linear regressions ran between *lack of self-control* PGS and WM MD within each ADI quantile. **C-D**) Dominance analysis illustrating the contribution of each variable (i.e., PGS or SES measure) to **C**) global brain measures at baseline or **D**) developmental change in global brain measures. The percent contribution of each input variable is calculatged as the variable s dominance relative to the total fit of the model. WM: White Matter, MD: Mean Diffusivity, ADI: Area Deprivation Index.

#### Dominance Analysis

We used dominance analysis to ascertain the relative contributions of genetics and environment to brain measures. Variance in mean cortical thickness and total subcortical volume was best captured by *lack of self-control* PGS (69.6% and 47.0%, respectively), while subcortical cellularity was best explained by *reward drive* PGS (HNI: 62.1%, RNI:35.3%). Notably, parental education contributed the most to total surface area (48.8%), and ADI and parental marital status were the biggest contributors for WM FA (24.9%) and WM MD (46.3%), respectively (Fig.8C).

Dominance analysis on longitudinal data showed that change of mean WM FA was best described by *lack of self-control* PGS (47.1%; Fig. 8D). ADI contributed most to change in average subcortical cellularity values (HNI: 42.0%, RNI:73.7%), as well as to change in WM MD (46.3%).

## Discussion

Using genomic SEM on 17 impulsivity and risk-taking traits, we identified three latent factors—which we labeled *lack of self-control*, *reward drive*, and *sensation seeking*—with both distinct and overlapping genetic associations. Furthermore, the polygenic scores for these factors were associated with specific brain morphology patterns during childhood. Finally, we demonstrated that socioeconomic factors can moderate the relationship between genetic predisposition and brain morphometry.

Our genomic SEM recapitulates the dual-process conceptualization of impulsivity, comprising “top-down” (poor cognitive control) and “bottom-up” (motivation) processes (28–32,119,120). Moreover, it further separates the “bottom-up” system into *reward drive* and *sensation seeking* factors. Gene enrichment analyses also revealed that genes implicated in *reward drive* were more preferentially involved in neurogenesis, whereas those associated with *sensation seeking* related to synaptic properties.

Consistent with this, *reward drive* was enriched for genes expressed during the prenatal period, when cortical neurogenesis occurs (121,122), while *sensation seeking* showed enrichment for genes expressed in late infancy, when synapse formation and pruning peak (123–126). Finally, *reward drive* PGS was mostly associated with subcortical nuclei, while *sensation seeking* PGS was reflected in cortical and white matter structure in children. These results collectively suggest that the motivational aspect of impulsivity can be separated into two components with distinct genetic and neurodevelopmental origins.

While the GWAS data used in our modeling were derived from adults, the genetic predisposition was already observable at the behavioral level in children. *Lack of self-control* PGS was indeed related to impulsivity in children—especially with UPPS-P subscales reflecting poor planning and high urgency. Similarly, *sensation seeking* PGS was associated with the UPPS-P Sensation Seeking scale itself in children. While manifestations of *reward drive* (drug use and sexual partners) are not present in children, its PGS was associated with externalizing behavior.

The three latent genetic factors showed distinct brain associations in children. *Lack of self-control* PGS was associated with reduced cortical thickness and surface area, most prominently in the frontal lobes. This aligns with previous findings linking lower self-control to reduced frontal gray matter across the lifespan (33,100,127–129), and supports the role of frontal lobes in self-regulation and executive function, as shown in functional neuroimaging (130,131) and lesion studies (132).

*Reward drive* PGS showed associations mainly with heightened cellularity of subcortical nuclei and reduced cortical thickness in orbitofrontal cortex. The subcortical and limbic structures associated with *reward drive* PGS, namely putamen, nucleus accumbens, amygdala, caudate, and orbitofrontal cortex are central to reward processing (133) and implicated in substance use (134–136). While a study in adults found reduced subcortical volumes to be associated with a reward-related phenotype (137), our study in children showed higher cellular density. As most children have not yet used addictive substances, this brain pattern may reflect a predisposing neurological state, whereas the reduction in volume observed in adults may be the result of substance use itself.

The *sensation seeking* PGS was associated with increased cortical surface area and higher white matter FA. This is consistent with evidence directly linking sensation seeking measures to cortical surface area and white matter integrity (103,138,139).

Cortical surface expansion and white matter myelination have been shown to co-occur in children, pointing to a shared neurodevelopmental origin that underpins cognitive functions (140). We also found that traits composing the *sensation seeking* factor exhibited positive genetic correlations with Educational Attainment. These results underscore *sensation seeking*’s unique nature and may suggest that this genetic factor is less likely to be associated with later negative outcomes.

Leveraging the longitudinal design of the ABCD Study, we found that *lack of self-control* PGS was associated with slower rate of increase in white matter FA, which likely captures myelination and axonal development (141). This association pattern differed from the cross-sectional results, suggesting dynamic genetic effects on brain development.

While the three factors showed distinct genetic and neurodevelopmental associations, they also showed similarities, particularly the implication *of CADM2* and *FOXP2*. *CADM2* encodes a synaptic cell adhesion protein and mediates synaptic plasticity. Previous work has shown its consistent association with impulsivity and risk-taking, as well as with a wide range of phenotypes such as anxiety and migraine (45). *FOXP2* encodes a transcription factor that regulates the embryonic expression of various genes, including those involved in neuronal growth and synaptic plasticity (127). This gene is highly expressed in cortex and basal ganglia (128) and plays a role in the synaptic wiring of the cortico-basal ganglia circuit (129), a critical network for the cognitive functions.

We also demonstrated the importance of socioeconomic factors in relation to genetic predisposition. Specifically, higher *lack of self-control* PGS was linked to reduced white matter integrity in children from more deprived environments, showing that the relation between genetic risk and the brain can vary according to environmental context. Furthermore, dominance analysis revealed different relative contributions of genetic and environmental factors in brain structure in children (59–62). For cortical thickness, subcortical volume, and subcortical cellularity, PGSs for the three factors had larger total contributions than the total of SES factors, whereas for cortical surface area and white matter integrity, SES factors contributed more than PGSs.

Our model differed from previous genomic SEM models on impulsivity, mainly due to the data-driven approach and the inclusion of risk-taking traits (55,142). For example, Gustavson et al. (55) constructed a 5-factor model that differentiates the UPPS-P subscales, also demonstrating that Delay Discounting constitutes a separate factor. Our model, on the other hand, grouped Lack of Premeditation, Negative and Positive Urgency, and Delay Discounting—which were genetically correlated—as the *lack of self-control* factor. Our data-driven procedure rather picked up the genetic overlap of these traits and highlighted the significant differences from the other traits included.

There are limitations in our study. First, our analyses were limited to individuals of European ancestry, limiting generalizability. This highlights the need to include more diverse populations in future studies (143). Next, the GWAS effective sample sizes differed across the latent factors, which made it difficult to compare results from gene-based analyses. Specifically, the *lack of self-control* factor had a smaller sample size, making it unclear whether the absence of overlap in the enrichment analyses truly represented the difference from the other two factors. Additionally, while our PLS analyses showed collective characteristics of the brain structure associated with each PGS, it remains unclear whether these reflect a unitary neurobiological influence or different independent developmental processes. Finally, this study only focused on brain-PGS associations that were observed across both sexes. How the effect differs between males and females should be explored in the future, as they may show divergent neurodevelopmental patterns.

In sum, this study highlighted the multifaceted nature of impulsivity and risk-taking at the genetic level. Furthermore, we show that the genetic predisposition for these traits is manifested in the brain during early development, with each facet involving different brain structures. This underscores that impulsivity and risk-taking comprise multiple dimensions that tap into different brain mechanisms, which are shaped by unique genetic and neurodevelopmental factors.

## Acknowledgements

This work was supported by funding from the Canadian Institutes of Health Research and Natural Sciences and Engineering Research Council of Canada to AD. MS was supported by the Funai Foundation for Information Technology fellowship and Fonds de recherche du Québec–Santé (FRQS) Doctoral fellowship. UV has been funded by Estonian Research Council’s personal research funding start-up grant PSG759.

Data used in the preparation of this article were obtained from the Adolescent Brain Cognitive Development^SM^ (ABCD) Study (https://abcdstudy.org), held in the NIMH Data Archive (NDA). This is a multisite, longitudinal study designed to recruit more than 10,000 children age 9-10 and follow them over 10 years into early adulthood. The ABCD Study® is supported by the National Institutes of Health and additional federal partners under award numbers U01DA041048, U01DA050989, U01DA051016, U01DA041022, U01DA051018, U01DA051037, U01DA050987, U01DA041174, U01DA041106, U01DA041117, U01DA041028, U01DA041134, U01DA050988, U01DA051039, U01DA041156, U01DA041025, U01DA041120, U01DA051038, U01DA041148, U01DA041093, U01DA041089, U24DA041123, U24DA041147. A full list of supporters is available at https://abcdstudy.org/federal-partners.html. A listing of participating sites and a complete listing of the study investigators can be found at https://abcdstudy.org/consortium_members/. ABCD consortium investigators designed and implemented the study and/or provided data but did not necessarily participate in the analysis or writing of this report. This manuscript reflects the views of the authors and may not reflect the opinions or views of the NIH or ABCD consortium investigators.

The ABCD data repository grows and changes over time. The ABCD data used in this report came from 10.15154/z563-zd24. DOIs can be found at http://dx.doi.org/10.15154/z563-zd24.

