## Supplementary Information for "Genomic structural equation modeling of impulsivity and risk-taking traits reveals three latent factors distinctly associated with brain structure and development"

***Meta-analysis of Risk Tolerance and Educational Attainment GWAS***

A meta-analysis of risk tolerance GWAS was performed using the standard error-based approach implemented in METAL (1), combining summary statistics from the discovery general risk tolerance GWAS in the UK Biobank and 10 replication cohorts (total n= 466,571) with those from the 23andMe GWAS dataset (n=969,298). Meta-analysis of Educational Attainment was also run with METAL, following the original paper by Okbay et al. (2)

***Linkage disequilibrium score regression***

We used the *LD Score* v1.0.1 to estimate genetic correlations (3). The package was also used to calculate LD Score intercepts for the multivariate GWAS results (4). We used the *scipy* package in Python to perform hierarchical clustering on the genetic correlation matrix.

***Genomic Structural Equation Modeling***

*Genomic SEM* R package v0.0.5 (5) was used to build and test factor models, and to perform multivariate GWAS. Alleles from different summary statistics were aligned using the HapMap3 reference panel. We removed SNPs with an INFO score below 0.9 or a minor allele frequency (MAF) under 0.01. The default diagonally weighted least squares estimation was used for the analyses.

Exploratory factor analysis

We first constructed a genetic covariance matrix using information only from even-number chromosomes, as exploratory factor analysis (EFA). Odd-number chromosomes were used for the confirmatory analysis step (below) to avoid overfitting of the model. The genetic covariance matrix was submitted to the *factanal* function from the *stats* R package v4.3.1 to perform EFA. We tested solutions with one to four latent factors.

Confirmatory factor analysis

Using parameters from EFA, we ran confirmatory factor analysis on odd-number chromosomes to establish model fit. Specifically, we constructed common-factor, two-factor, three-factor, and four-factor models by retaining indicators that had standardized loadings higher than 0.5 in EFA results (This process dropped indicators that did not have ≥0.5 loadings to any factor. For example, EA, BMI, and UPPS-P Lack of Perseverance indicators were not modeled in the three-factor model). Model fit was examined using the comparative fit index (CFI) and the standardized root mean square residual (SRMR). The model fit indices for each model were: common-factor model CFI=0.686, SRMR=0.156; two-factor model CFI=0.849, SRMR=0.126; three-factor model CFI=0.865, SRMR=0.104; four-factor model CFI=0.813, SRMR-0.099.

Additionally, we added residual covariances between variables to address possible shared liability that are not modeled with the latent factors; residual covariances between UPPS-P Sensation Seeking and other UPPS-P subscales, BIS subscales, and Delay Discounting were added to account for the shared disposition. We also added residual covariances between Risk Tolerance and BIS Motor, UPPS-P Lack of Premeditation, and UPPS-P Positive Urgency due to the similarity in specific items asked in the questionnaires.

Multivariate GWAS

We omitted SNPs that had $Q_{SNP}$ p-values below 5e-8. $Q_{SNP}$ is a test statistic that measures how much a given SNP operates on the individual phenotypes exclusively through the latent factor. A larger $Q_{SNP}$ value (i.e., significant p-value) indicates that the effect of SNP on the individual phenotypes is not fully captured through the latent model.

The SNP2GENE function from the Functional Mapping and Annotation (FUMA) v1.5.2 pipeline (6) was used to identify genomic risk loci, lead SNPs, and all independent significant SNPs from each latent factor identified by genomic SEM. Briefly, FUMA identifies lead SNPs through a double-clumping procedure. The independent significant SNPs were first found by clumping SNPs with *p*-value < 5e-8 and independent at $r^{2}\leq$ 0.6. The lead SNPs were defined by applying a second round of clumping with *p*-value < 5e-8 and $r^{2}\leq$ 0.1. Finally, genomic risk loci are defined by grouping independent significant SNPs that are dependent with each other ($r^{2}$ ≥ 0.1) or close to each other (250 kb).

We observed inflation in test statistics for all factors (*lack of self-control*: mean χ^2^ = 1.20, λ_GC_ = 1.17; *reward drive*: mean χ^2^ = 2.13, λ_GC_ = 1.72; *sensation seeking*: mean χ^2^ = 2.05, λ_GC_ = 1.70), suggesting polygenicity (Fig. S3 for QQ plots). The LDSC regression intercepts further indicated that the inflation was indeed due to the polygenic structure rather than population stratification (*lack of self-control* LDSC intercept=0.99, SE=0.007; *reward drive* intercept=1.00, SE=0.01; *sensation seeking* intercept =1.07, SE= 0.01).

***Genetic Correlation with Brain Characteristics***

We further asked whether there were overlaps in the genetic architecture between brain structures and *lack of self-control, reward drive,* and *sensation seeking* factors. We conducted genetic correlation analyses between the latent factors and brain structural measurements using brain imaging GWAS data from UK Biobank (7). Specifically, we looked into GWASs of cortical surface area and thickness (DK atlas), volumes of subcortical structure, and FA and MD of WM tracts from Probtrack atlas (8–10). We found no significant association after multiple comparison corrections (Fig. S4).

***Gene-based analyses***

We used MAGMA v1.08 (11) to conduct gene association, gene-set enrichment, and gene property analyses for each latent factor. Default MAGMA parameters were used throughout the analyses. The SNPs were mapped to 19,155 protein coding genes from Ensembl v102. For gene set enrichment analyses, 15,496 gene sets (curated gene sets: 5,500, GO terms: 9,996) from MsigDB v7.0 were tested. For gene property analyses, 54 tissues from the GTEx v8 were tested. Additionally, we used brain tissue data from BrainSpan dataset to test 11 developmental epochs. The significance was determined based on the Bonferroni-corrected thresholds; $P\leq3.22\times{10}^{-6}$ for gene sets, $P\leq9.26\times{10}^{-4}$ for tissues, and $P\leq4.54\times{10}^{-3}$ for BrainSpan analyses.

***Quality Control of the Genetic Data in the ABCD dataset***

Quality control for both samples and variants was performed following standard guidelines (<https://github.com/neurogenetics/GWAS-pipeline>) using *plink* 1.9. We used GCTA to check for relatedness and excluded individuals who have a first or second degree relative in the cohort (π>0.125). Imputation was performed on the Michigan Imputation Server using reference panel HRC 1.1 2016 and Eagle v2.3 Phasing.

***Partial Least Square***

Partial least squares (PLS) analysis was used to relate genetic predispositions and brain structures. PLS is a multivariate method that decomposes the two input datasets and finds latent factors, in which weighted linear combinations of original variables maximally covary with each other. In our case, the two matrices were data of PGSs (4124 subjects $\times$ 1 PGS) and brain features (4124 subjects $\times$ 234 brain features). This allowed us to identify patterns of brain characteristics that were associated with the PGS. Both matrices were z-scored. Additionally, we accounted for covariates, i.e., age at baseline, sex, 15 genetic principal components (PCs), and genetic batch by regressing them out from the brain measures as covariates.

PLS first computes the covariance matrix of the two matrices, which represents the covariation of the PGS and brain measures across the participants. This matrix was then subjected to singular value decomposition, which finds the latent factors that effectively associate PGS and brain features. This consists of weights for original variables as well as singular values; the weights represent how much each variable contributes to the latent factors, and the singular values are proportional to the total PGS-brain covariance accounted for by the latent factor. We then projected the original data onto the latent space using the respective weights, yielding a brain score for each individual.

The significance of each latent factor was assessed using a permutation test. We randomly permuted the ordering of participants in the brain measurement matrix, and recalculated the covariance matrix between PGSs (unpermuted) and brain measures (permuted). This covariance matrix was then subjected to singular vector decomposition, generating a singular value under the null hypothesis that there is no relation between PGSs and brain measures. This procedure was repeated 500 times to generate a null distribution, against which the original singular value was tested.

Bootstrap resampling was used to assess the contribution of individual variables (PGSs or brain measures) to each latent factor. The rows (participants) of both matrices were randomly selected with replacement and a resampled covariance matrix was calculated. This was then subjected to singular vector decomposition. By repeating this procedure (N=500), we obtained a sampling distribution for each individual weight. A bootstrap ratio (BSR) was calculated for each variable as the ratio of the original weight and its bootstrap-estimated error. Thus, variables with large bootstrap ratios had larger weights (i.e., high contribution to the latent factor) with small standard errors (i.e., high reliability). Since BSR is a proxy of z-score, BSR values >1.96 or <−1.96 are equivalent to having *p* < 0.05 significance.

We tested the robustness of the PLS models by cross-validating the correlation between PGS and brain scores. Following previous studies (12,13), we reran the analyses with 100 randomized train and test splits, where test set is 25% of the original dataset. PLS models were constructed with the training set and projected on the test set to derive predicted brain scores. These brain scores were then correlated with respective PGSs. The resulting correlation coefficient was tested against permutation null distribution to assess significance.

Directionality of PLS associations

Given the inherent indeterminacy of association direction in PLS analyses, we used brain scores to capture the covariation direction between PGSs and brain measures. Brain score reflects how well a participant express the latent brain pattern, obtained by projecting the latent brain pattern weights onto the original brain characteristic matrix. For cross-sectional analyses, all PGSs had a positive association with their corresponding brain scores (r(4117) = 0.091 for *lack of self-control* PGS; r(4117) = 0.092 for *reward drive* PGS; r(4117) = 0.088 for *sensation seeking* PGS; Fig. S5a-c). This allowed us to directly translate the BSR to the association direction, i.e., if BSR was positive, the association between the PGS and brain measure was positive, whereas negative BSR indicated a negative association.

For longitudinal analysis, we also observed a positive association between *lack of self-control* PGS and corresponding brain score (Fig. S5d). This allowed us to directly use BSR to obtain association direction. To further contextualize, we translated the BSR to the level of development (Fig. 5b). For example, for a measurement whose general developmental trajectory is increasing, having a positive BSR indicates that a higher *lack of self-control* PGS is related to a bigger change (accelerated development), while having a negative BSR represents smaller change (slowed development).

***Dominance Analysis***

Dominance analysis is a statistical technique used to determine the relative importance of each predictor in explaining the variance in the target variable. This is done by building regression models with all possible subsets of predictors and assessing which one consisitently improves the model’s predictive power (i.e., R^2^) in every combination of variables. In our study, we examined if the variance in each global measurement was better explained by the genetic predisposition or socioeconomic factors.

**Supplementary Figures**

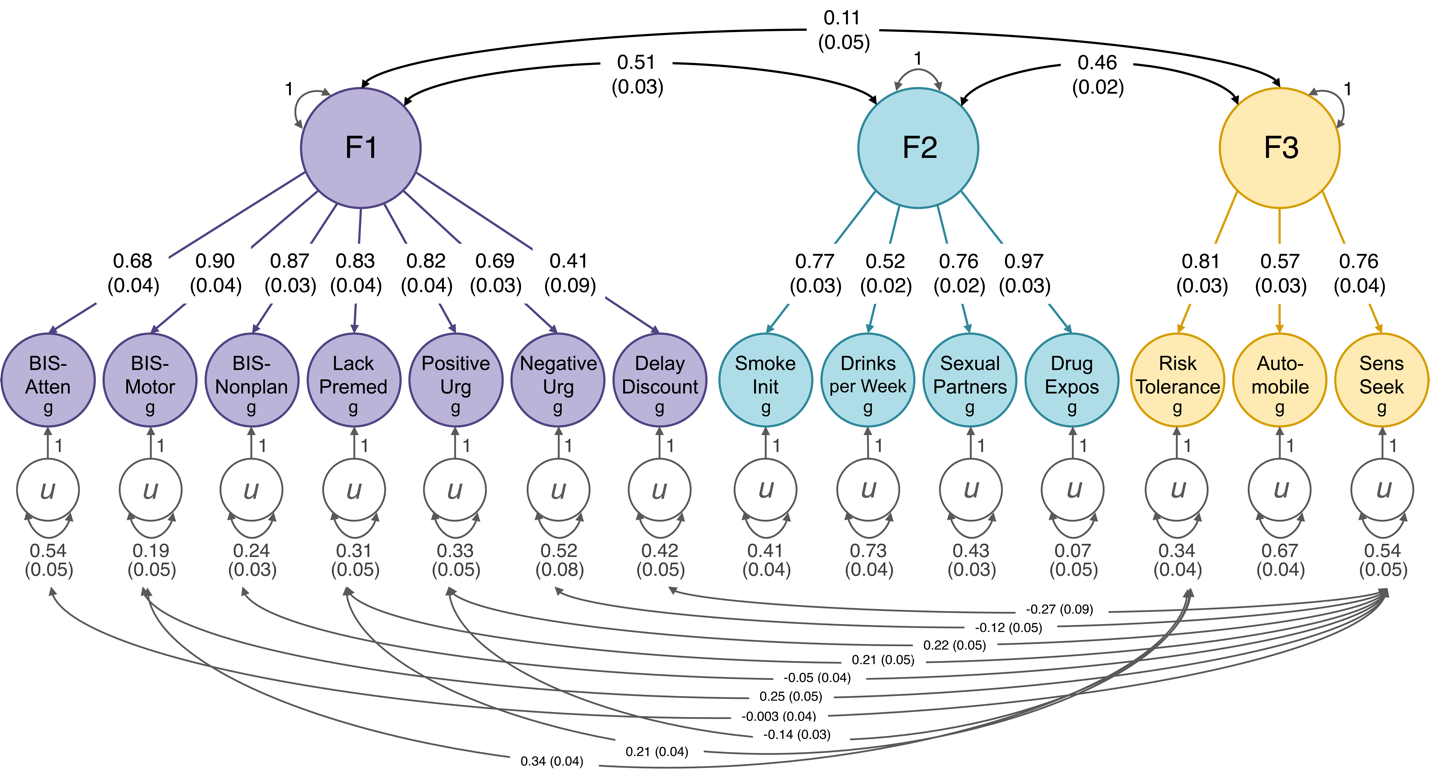

**Figure S1**

Residual covariances were included to account for potential shared genetic liability. Since UPPS-P Sensation Seeking is administered alongside other UPPS-P subscales, which constitute the majority of F1 (*lack of self-control*), covariances were added between these indicators. Additionally, BIS-Motor, UPPS-P Lack of Premeditation, and UPPS-P Positive Urgency include questionnaire items with conceptual overlap with the Risk Tolerance questions (e.g. Are you comfortable with taking risks?), warranting the inclusion of residual covariances (e.g. “I spend or charge more than I earn.” in BIS-Motor, “I have a reserved and cautious attitude toward life” in UPPSP Lack of Premeditation, “Others are shocked or worried about the things I do when I am feeling very excited.” in UPPSP Positive Urgency.) F1: lack of self-control, F2: reward-drive, F3: sensatin seeking.

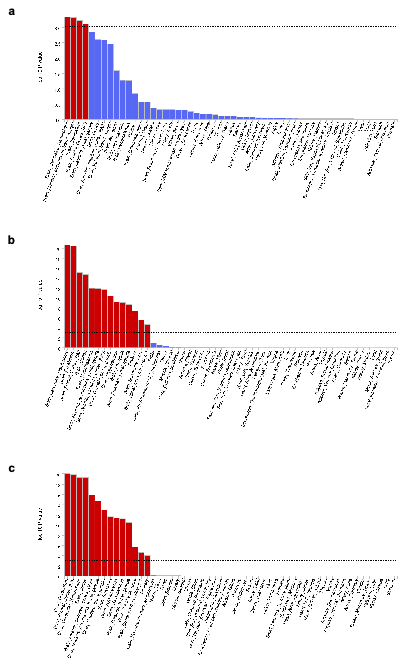

**Figure S2**

MAGMA tissue expression analysis results. Per-tissue enrichment of expression of genes associated with **a**) *lack of self-control*, **b**) *reward drive*, or **c**) *sensation seeking*, based on GTEx RNA-seq data for 53 specific tissue type.

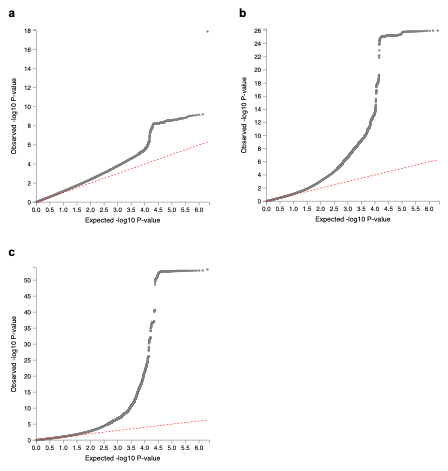

**Figure S3**

QQ plots from multivariate GWAS of a) *lack of self-control*, b) *reward drive*, and c) *sensation seeking*, showing the degree of inflation of all test statistics.

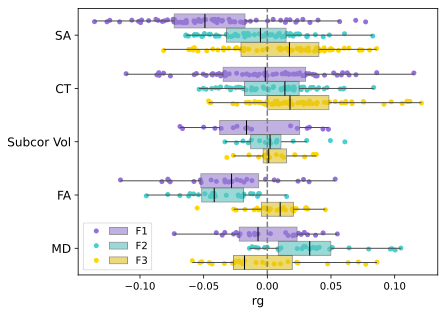

**Figure S4**
Genetic Correlation between the latent factors and brain structural measures. Brain imaging GWAS data from UK Biobank data. Each dot represents genetic correlation coefficient between a brain structure measure and the impulsivity/risk-taking latent factor. F1: lack of self-control, F2: reward drive, F3: sensation seeking, SA: cortical surface area, CT: cortical thickness, FA: white matter fractional anisotropy, MD: white matter mean diffusivity.

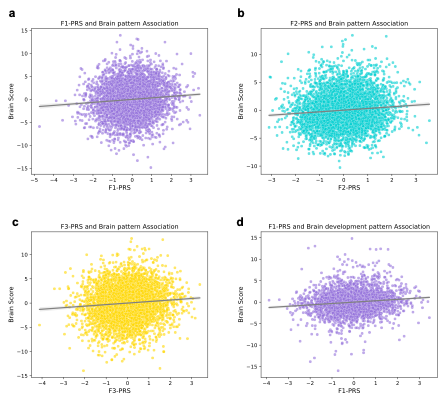

**Figure S3 Association between brain score and PGS**

**a-c**) Scatther plot of brain composite scores **a**) lack of self-control PGS, **b**) *reward drive* PGS, and **c**) *sensation seeking* PGS. Brain composite scores are derived from each PLS analysis with baseline data. The positive association indicates that the brain weights (Bootstrap Ratio) in Fig. 4 directly reflect the direction of association with the corresponding PGS—positive BSR indicates a positive association, while negative BSR indicates a negative association. **d**) Scatter plot of brain composite score from PLS based on brain developmental data and *lack of self-control* PGS.
